# Blimp-1 shapes germinal center B cell clonal diversity by gating chromatin accessibility during light-to-dark zone transition

**DOI:** 10.1101/2025.08.27.672737

**Authors:** Godhev Kumar Manakkat Vijay, Bala Ramaswami, Steven Gierlack, Nicholas A. Pease, Peter Gerges, Dianyu Chen, Kairavee Thakkar, Luis Mena Hernandez, Swapnil Keshari, Heping Xu, Nathan Salomonis, Jishnu Das, David M. Rothstein, Harinder Singh

## Abstract

Germinal center (GC) B cell responses are defined by many positive regulators of affinity maturation, but few components that restrain clonal dominance, notably *Nr4a1*, are known. We reveal an unsuspected role for Blimp-1 (*Prdm1*), a plasma cell determinant, as a feedback regulator of affinity maturation. Single cell RNA-seq and B cell receptor (BCR)-seq showed that B cell specific *Prdm1* loss drives an exaggerated GC reaction with larger clones, increased somatic hypermutation, and greater clonal dominance, independent of *Nr4a1*. Single cell chromatin profiling with base-resolution modeling indicated that Blimp-1 represses expression of BCR-signaling genes, gating chromatin accessibility at ISRE, EICE, NF-κB, and Oct motifs. In the absence of Blimp-1, enhanced BCR signaling augments activities of transcription factors that promote G1–S transition during light-zone (LZ) selection and fuel dark-zone (DZ) expansion. Thus, Blimp-1 attenuates BCR signaling and constrains the LZ to DZ transition, fine-tuning clonal competition thereby maintaining repertoire diversity.

## Introduction

Extensive studies have delineated signaling systems and transcription factors (TFs) that positively regulate GC B cell responses^1–11^. In the LZ, T follicular helper (T_fh_)-derived CD40L and ICOS along with B cell receptor (BCR) signals license B cell clones to pass through the G1-S checkpoint; the cells then re-enter the DZ for clonal expansion and AID-mediated somatic hypermutation (SHM). Among TFs that promote GC initiation, selection, and output, BCL6 programs GC identity. MYC licenses positively selected LZ cells for re-entry and proliferation in the DZ, whereas FOXO1 controls the program underlying rapid cell divisions. PU.1 and SPIB via interactions with IRF8 help maintain GC identity and selection, acting on ETS–IRF composite elements (EICE)^12,13^. BATF supports robust GC responses and class switching^14^. The POU TF module comprised of OCT1/POU2F1, OCT2/POU2F2, and their co-activator OBF1/POU2AF1 maintains BCR-dependent transcription required for GC expansion^15^. Canonical and alternative NF-κB components c-Rel and p52/RelB also promote GC maintenance. A p52–ETS1 complex has been recently shown to induce OCT1 and OBF1, thereby generating a feedforward regulatory loop^16,17^.

In contrast, few signaling-induced negative regulators of the GC response that restrict clonal dominance have been identified, notably *Nr4a1* (NUR77)^18,19^. We posited that Blimp-1 which promotes exit of GC B cells into the plasma cell (PC) pathway, could also act as a signaling-induced transient feedback gate to restrain the GC B cell response, thereby reducing clonal dominance. In keeping with this hypothesis, low and heterogenous expression of Blimp-1 has been reported in GC B cells and its loss has been shown to result in larger GCs^20,21^. Furthermore, Conter et al. recently demonstrated using bone marrow chimeras that B-cell-intrinsic Blimp-1 loss results in more exuberant GC responses associated with enhanced proliferation of antigen-specific GC B cells^22^. We note that in addition to the feedback gating model for Blimp-1 action in the GC considered by us, two other models could account for the GC phenotype: (i) increased GC seeding due to impaired plasmablast differentiation or (ii) failed GC exit due to blocked PC differentiation. We reasoned that integrated single cell (sc) transcriptional and chromatin analyses of GC B cells along with delineation of their clonal dynamics could distinguish among the three models. Furthermore, such high dimensional analyses of the perturbed and amplified GC response would yield insights into the underlying molecular mechanisms of Blimp-1 action.

Using sc transcriptome, V(D)J and chromatin profiling coupled with single nucleotide-resolution accessibility modeling of regulatory DNA sequences, we show that *Prdm1*-deficient B cells mount an exaggerated GC reaction characterized by larger clone sizes and enhanced affinity maturation culminating in greater clonal dominance. This phenotype is not attributable to increased GC seeding nor can it be merely accounted for by impaired exit of GC B cells into the PC pathway. Rather, the integrated genomic analyses reveal that Blimp-1 constrains expression of genes encoding components of the BCR-signaling cascade. Loss of this feedback, results in enhanced *Syk*, *Lyn* and *Btk* activity in GC B cells and augments chromatin engagement of signaling-inducible TFs at EICE, NF-κB and POU (OCT) motifs, promoting the G1–S transition during LZ selection and fueling DZ expansion.

## Results

### Blimp-1 deficient B cells undergo more robust GC response

To test the hypothesized function of Blimp-1 in GC B cells, we utilized *Prdm1*^fl/fl^*Cd19*^cre/+^ (*Prdm1* cKO) and *Prdm1*^+/+^*Cd19*^cre/+^ (WT) mice and analyzed their GC responses under steady state conditions and after immunization with NP-CGG alum and LPS (Fig. 1a). Loss of Blimp-1 resulted in increased numbers of GC B cells at steady state and at 7- and 14-days post-immunization (d.p.i.) with NP-CGG (Extended data Fig. 1a). Blimp-1 deficiency also led to increased numbers and frequencies of CD80^+^ and PD-L2^+^ memory B cells^23^ at both 7 and 14 d.p.i. (Extended data Fig. 1b,c), consistent with the expanded GC response and suggesting Blimp-1 deficiency also enhances post-GC differentiation toward the memory fate. As anticipated, frequencies and numbers of NP^+^ GC B cells were increased in *Prdm1* cKO mice at 7 and 14 d.p.i. (Fig. 1b,c and Extended data Fig. 1d,e).

**Fig. 1.**
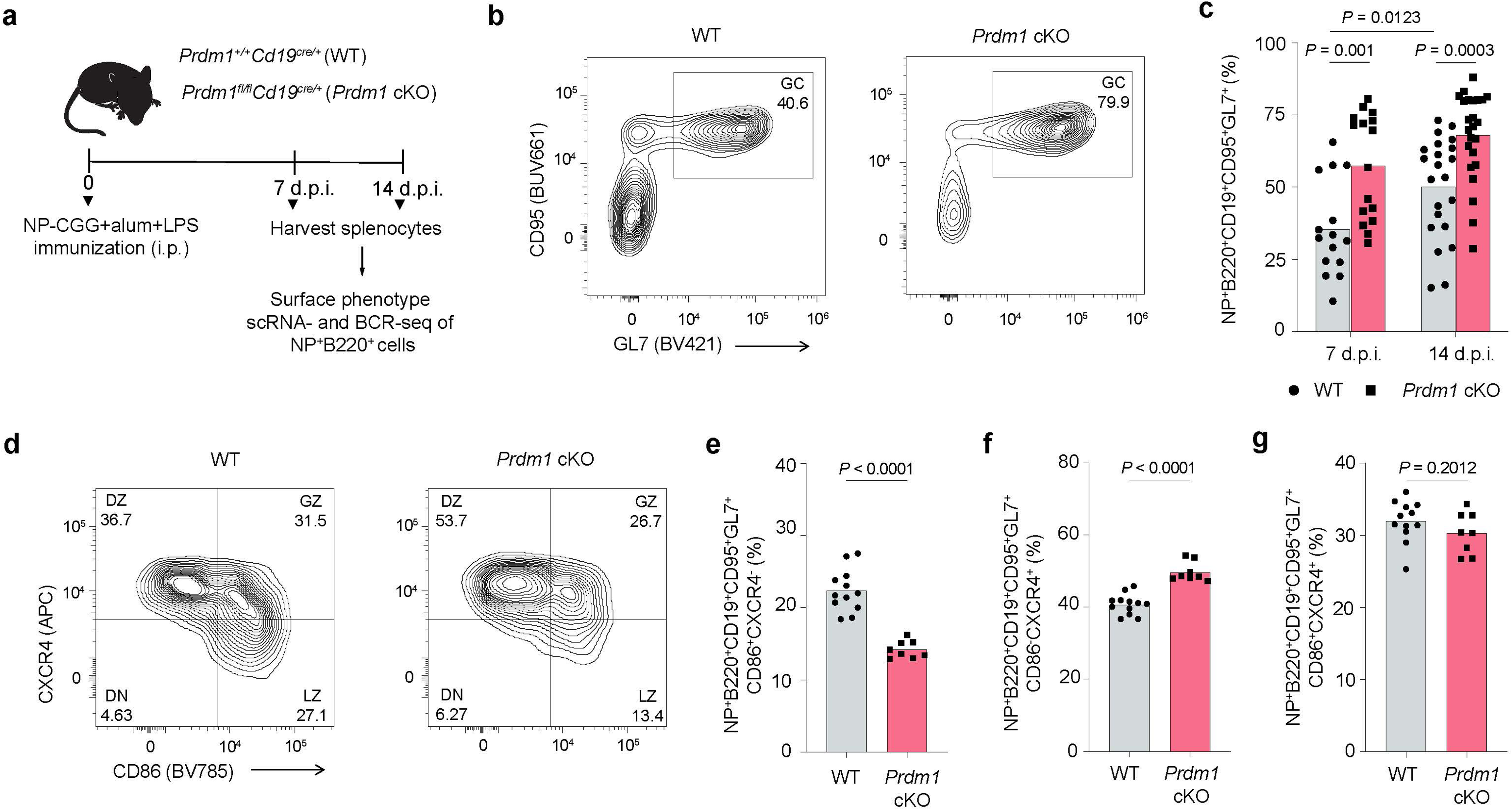
Blimp-1 loss results in GC expansion and positively skews DZ/LZ ratio. **a,** Experimental design for analyzing frequencies, single cell transcriptional states and BCR sequences of NP-specific B cells elicited by immunization of WT and *Prdm1* cKO mice. **b,** Representative flow plots showing the frequency of NP^+^ GC B cells in WT and *Prdm1* cKO mice (14 d.p.i.). **c,** Frequencies of NP^+^ GC B cells in WT (n=15) and *Prdm1* cKO (n=16) mice at 7 d.p.i. and WT (n=22) and *Prdm1* cKO (n=24) mice at 14 d.p.i. **d,** Representative flow cytometry plots showing the frequencies of LZ, GZ and DZ GC B cells in WT and *Prdm1* cKO mice. **e-g,** Frequencies of NP-specific LZ, DZ and GZ GC B cells in WT (n=12) and *Prdm1* cKO (n=8) mice (14 d.p.i.) respectively. Each symbol (**c, e-g**) represents an individual mouse. Data in panels **b,c** are from three different experiments and **d-g** are from two independent experiments. Statistical significance was tested by two-tailed Mann-Whitney test (**c, e-g**).

To determine whether this overall expansion was accompanied by changes in the cellular composition of GC compartments, we examined the distribution of LZ, DZ and GZ (gray zone)^24^ GCBCs. Although absolute numbers of LZ, DZ and GZ GCBCs were increased in the *Prdm1* cKO mice, this was accompanied by a decreased relative frequency of LZ GCBCs and increased frequency of DZ GCBCs. The frequency of the GZ population remained unchanged (Fig. 1d-g and Extended Data. Fig. 1f-h). To confirm B-cell-intrinsic functions of Blimp-1, we generated bone marrow chimeras by reconstituting irradiated recipients with WT and *Prdm1* cKO bone marrow. The GC expansion phenotype was reproduced, demonstrating that Blimp-1 constrains the GC response in a B-cell-intrinsic manner (Extended Data Fig. 1i,j). These results were consistent with the hypothesis that Blimp-1 constrains the GC response, during the LZ to DZ transition.

### scRNA-seq reveals Blimp-1 constrains LZ to DZ transition

To analyze the impact of Blimp-1 perturbation on transcriptional programming during the LZ to DZ transition, we performed scRNA-seq on NP-specific B cells^7^ at 14 d.p.i. Analyses of the WT scRNA-seq datasets using the ICGS2 computational pipeline^25^ identified five distinct cell clusters that could be distinguished based on biologically informative, marker genes (Fig. 2a, Extended Data Fig. 2a and Supplementary Table 1). Within the two LZ clusters, LZ-1 manifested the highest expression of B cell activation markers, including *Gpr183*, *Cd55* and *Cd38*. In contrast, LZ-2 showed elevated expression of genes involved in antigen presentation and BCR signaling, including *Cd83*, *H2-DMa*, *H2-Oa* and *Nfkbia*. The two DZ clusters, DZ-1 and DZ-2, resolved cells within the G1-S and G2-M phases of the cell cycle, respectively. DZ-1 was characterized by high expression of DNA replication genes (*Mcm5*, *Mcm7*, *Pcna)*, whereas DZ-2 manifested higher expression of mitotic components and their regulators, including *Cenpf*, *Ccnb1* and *Cdk1* (Fig. 2a and Supplementary Table 2). Notably, the analysis identified a distinctive GCBC cluster, herein termed transition zone (TZ), which exhibited elevated expression of ribosomal biogenesis and translation genes, including *Npm1*, *Nop58*, *Fbl* and *Eif5a* (Fig. 2a,b and Supplementary Table 2). Consistent with this transcriptional module, TZ cells showed the highest expression of *Myc* target genes, highlighting their importance as key intermediates that integrate activation and proliferation cues during the transitioning of LZ cells into the DZ^26,27^ (Fig. 2c). Gene signature analyses using curated GC B cell gene sets^7^ substantiated the conclusion that the TZ cluster represents an intermediate state, as the cells displayed the highest *Myc* score and also an increase in G1/S associated genes compared with LZ cells (Extended Data Fig. 2b-d). We note that a subset of TZ cells also expressed increased levels of transcripts that have previously been associated with cells transitioning from the LZ to the DZ, termed GZ GCBC^24^ (Fig. 2d, Extended Data Fig. 2e,f and Supplementary Table 3). To substantiate the developmental trajectory inferred by the annotations of the ICGS2 clusters based on their marker genes, we performed RNA velocity using scVelo^28^, which leveraged the dynamics of spliced and unspliced transcripts. This analysis supported the inference that TZ cells arise from LZ-2 and give rise to DZ-1 cells (Fig. 2e). We additionally performed analyses of the scRNAseq dataset using Monocle2^29^, which reinforced the inference that TZ cells reflect an intermediate state between LZ and DZ compartments (Extended Data Fig. 2g).

**Fig. 2.**
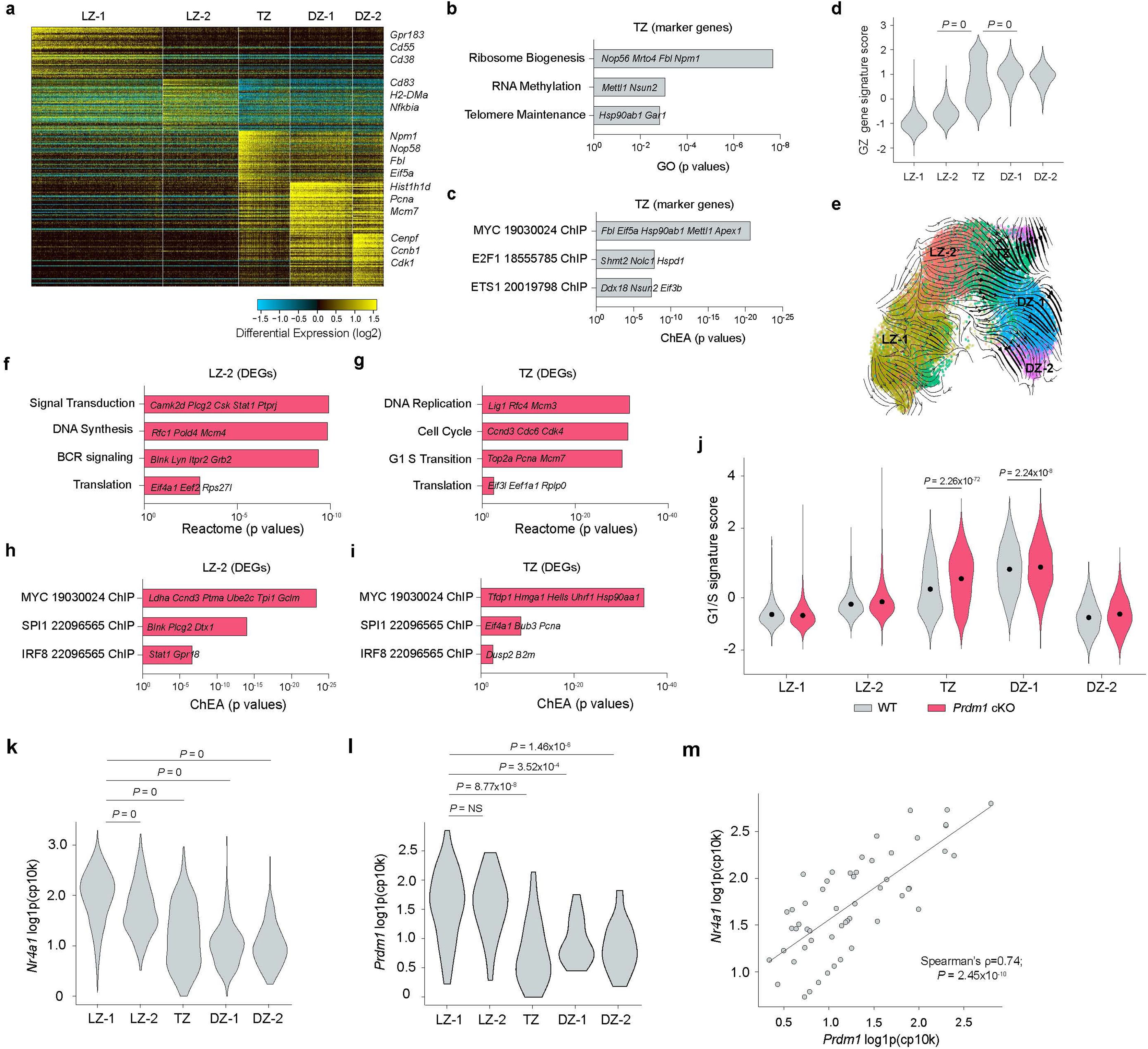
Blimp-1 constrains the GC B cell response by antagonizing the LZ to DZ transition. **a,** scRNA-seq analysis of splenic NP^+^B220^+^ cells isolated at 14 d.p.i. from WT mice (n=3). Heatmap was generated using cluster-specific marker genes delineated by the MarkerFinder algorithm in AltAnalyze **(Methods).** Columns in heatmap represent cells (n=28,809); rows represent MarkerFinder genes (n=300). Select MarkerFinder genes for each cell cluster are displayed on right of Heatmap. Cell clusters were generated using ICGS2 within AltAnalyze and annotated with their inferred genomic states. LZ, TZ and DZ refer to light zone, transitional zone and dark zone GC B cell states, respectively. **b,c,** Bar plots displaying key GO pathways **(b)** and TF target genes from the ChEA datasets **(c)** associated with the top 60 genes in TZ state. **d,** Violin plot displaying the GZ signature scores for the indicated cell clusters. **e,** Developmental trajectory of GC B cells constructed by scVelo on 2-D UMAP space **(Methods). f-i,** Bar plots displaying key Reactome pathways and TF target genes from the ChEA datasets associated with the upregulated DEGs (*Prdm1* cKO versus WT) in LZ-2 and TZ states. **j,** Plot displaying the G1/S signature scores across indicated cell clusters in WT and *Prdm1* cKO mice (14 d.p.i., n=3) **(Methods)**. **k,l,** Violin plots displaying log1p(cp10k)-normalized expression of *Nr4a1* **(k)** and *Prdm1* **(l)** in non-zero expressing cells across the indicated cell clusters. **m,** Scatter plot displaying *Prdm1* versus *Nr4a1* expression in non-zero expressing cells. Spearman’s correlation and two-tailed p values are indicated. Each dot (**e**) represents an individual cell. Selected genes within pathways are displayed (**b, c, f-i**). Data in panels **a-m** are from three different experiments. Statistical significance was tested by one-tailed Fisher’s Exact test and Benjamini-Hochberg method for correction of multiple-hypotheses testing (**b, c, f-i**), two-tailed Mann-Whitney test (**j**) and two-tailed Kruskal-Wallis with Dunn’s multiple comparison test (**d, k, l**).

Next, cellHarmony was used to align the *Prdm1* cKO GC B cells with their WT counterparts^30^(Extended Data Fig. 2h). Consistent with the phenotypic analysis, the relative frequency of *Prdm1* cKO DZ GC B cells was increased, in relation to their LZ counterparts (Extended Data Fig. 2i). Furthermore, the scRNAseq analysis suggested that the inversion of LZ and DZ frequencies may occur within TZ cells. To explore the molecular mechanisms by which Blimp-1 impacts GC state transitions, we performed differential gene expression analyses. In keeping with the dominant molecular actions of Blimp-1 as a transcriptional repressor we observed a larger number of upregulated compared with downregulated differentially expressed genes (DEGs), 749 and 260, respectively (Supplementary Table 4). The largest number of upregulated DEGs were observed in the LZ-2 and TZ clusters (Extended Data Fig. 2j). In the LZ-2 cluster, *Prdm1* cKO GC B cells exhibited enrichment of genes involved in BCR signaling and translation (Fig. 2f and Supplementary Table 5). In contrast, genes associated with the G1/S transition, DNA replication and cell cycle progression were upregulated in the TZ cluster (Fig. 2g and Supplementary Table 5). Notably, both clusters showed increased expression of genes known to be regulated by the TFs MYC, SPI1 (PU.1) and IRF8 (Fig. 2h,i and Supplementary Table 5). Consistent with the above analyses, the G1/S signature scores were increased in TZ and DZ-1 clusters (Fig. 2j). Taken together these transcriptional alterations agreed with the hypothesized function of Blimp-1 as a negative feedback regulator, the absence of which in the LZ, results in enhanced BCR signaling and precocious programming of *Myc*-mediated protein translation and the G1-S transition.

Expression of Blimp-1 in GC B cells was previously reported as low and sparse^31,32^. To determine whether *Prdm1* expression is in accord with its proposed function in constraining the LZ to DZ transition, we quantified the expression of *Prdm1* across various WT GC clusters in comparison with *Nr4a1*. In keeping with its induction during B cell activation as a consequence of BCR signaling^18^, *Nr4a1* transcripts were detected with the highest frequency (∼30%) and median values, in the LZ-1 cluster (Fig. 2k and Extended data Fig. 2k). We note that *Nr4a1* expression was not impacted in *Prdm1* cKO GC B cells (Supplementary Table 4). Though *Prdm1* transcripts were only detected in a small fraction of WT GC B cells (∼1%), like *Nr4a1,* they manifested highest median values in the LZ-1 cells (Fig. 2l and Extended data Fig. 2l). Importantly, *Prdm1* and *Nr4a1* transcripts were strongly correlated in cells that displayed their coincident expression (Fig. 2m). Thus, despite sparse detection of *Prdm1* transcripts in LZ cells, its expression pattern was consistent with its induction in BCR activated LZ cells and in restraining the LZ to DZ transition not dependent on controlling *Nr4a1* expression.

### Blimp-1 attenuates affinity maturation and clonal dominance

The above results implied that affinity maturation and clonal dominance in the GC may be accelerated with the loss of Blimp-1. To directly test this possibility, we leveraged scBCR-seq data generated in parallel with the scRNA-seq. To analyze NP-specific GC B cell clones, we focused on those harboring V_H_186.2 heavy chain gene segment (IGHV1-72*01) rearrangements which dominate the NP-specific GC B cell response at its peak^7,32^ (Supplementary Table 6). The W33L amino acid substitution has been shown to increase NP-binding affinity of the BCR containing the IGHV1-72*01 gene segment by ten-fold and represents the dominant mutation after positive selection during a peak GC response^7^. In keeping with the increased numbers and relative frequencies of NP-specific DZ GC B cells, we noted larger clone sizes and increased SHM within *Prdm1* cKO GC B cells during the initiation of the GC B cell response (7 d.p.i.). However, enhanced affinity maturation was not observable at this early stage (Fig. 3a-c). As was the case at 7 d.p.i., the clonal sizes of the *Prdm1* cKO GC B cells were significantly larger (14 d.p.i.) than their WT counterparts (Fig. 3d) suggesting that mutant cells in the DZ are expanded not only due to more efficient LZ to DZ transition but possibly due to faster cell cycle times. In keeping with these observations, the mutation frequencies of the IGHV1-72*01 gene segment were also increased in *Prdm1* cKO GC B cells at 14 d.p.i. (Fig. 3e). Importantly at the peak of the GC response, there was a significant increase in the frequency of W33L mutation bearing clones in the absence of Blimp-1 (Fig. 3f).

**Fig. 3.**
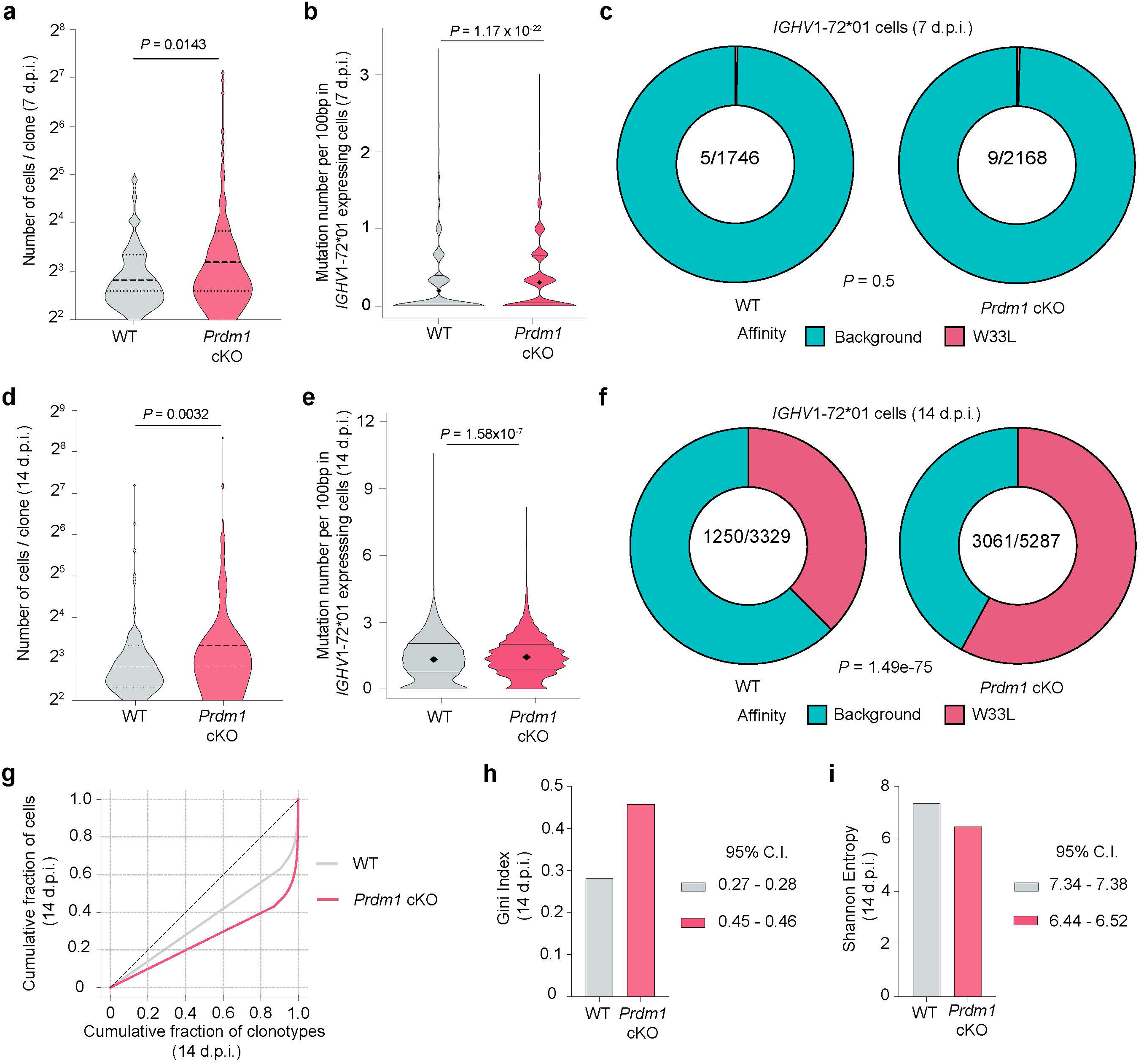
Blimp-1 constrains affinity maturation and clonal dominance. **a,** scBCR-seq analysis for the numbers of cells in distinct *IGHV1-72*01* clones in WT and *Prdm1* cKO mice (7 d.p.i.). **b,** scBCR-seq analysis for the mutation frequencies within *IGHV1-72*01* gene segments of GC B cells in WT and *Prdm1* cKO mice (7 d.p.i.). **c,** Pie-charts displaying the proportion of *IGHV1-72*01* gene segments harboring high affinity W33L mutations in splenic NP^+^B220^+^ cells of indicated genotypes (7 d.p.i.). **d,** scBCR-seq analysis for the numbers of cells in distinct *IGHV1-72*01* clones in WT and *Prdm1* cKO mice (14 d.p.i.). **e,** scBCR-seq analysis for the mutation frequencies within *IGHV1-72*01* gene segments of GC B cells in WT and *Prdm1* cKO mice (14 d.p.i.). **f,** Pie-charts displaying the proportion of *IGHV1-72*01* gene segments harboring high affinity W33L mutations in splenic NP^+^B220^+^ cells of indicated genotypes (14 d.p.i.). **g-i,** Clonal diversity of BCR sequences in WT and *Prdm1* cKO GC B cells (14 d.p.i.). **g,** Lorenz plots displaying the clonal distribution, skewed curves reflect clonal diversity. **h,i,** Bar plots displaying the Gini Index values **(h)** and the Shannon Entropy scores **(i)** for the BCR repertoires, respectively. Samples were collected from WT (n=3) and *Prdm1* cKO (n=3) mice at 14 d.p.i. and WT (n=1) and *Prdm1* cKO (n=1) mice at 7 d.p.i. For comparison across samples in h-i, data was rarefied to a common paired-cell depth per sample, computed with 95% confidence intervals and represented as mean values. Data in panels **d-i** are from three different experiments. Statistical significance was tested by two-tailed Mann-Whitney test (**a,b,d,e**) and Chi square test (**c,f**).

To analyze clonal dominance, we enumerated all clonotypes based on their paired IgH and IgL sequences in WT or *Prdm1* cKO NP-specific B cells. We then constructed Lorenz curves that display cumulative fraction of cells versus cumulative fraction of clonotypes, after ranking clonotypes from smallest to largest. In such an analysis a repertoire with every clonotype being equally represented, would generate a perfectly uniform distribution represented by a diagonal (Fig. 3g and Supplementary Table 7). As expected for non-uniform Ig repertoires, both the WT and *Prdm1* cKO distributions were displaced from the diagonal. Importantly, the *Prdm1* cKO distribution showed greater skewing from the diagonal indicating that a smaller subset of clonotypes were accounting for a large fraction of cells (14 d.p.i.). Next, we used the Gini coefficient to quantify this skewing based on area defined by the observed Lorenz curve and the diagonal. Values of the Gini coefficient can range from 0 (uniform repertoire) to 1 (maximal clonal dominance with all cells being of a single clonotype). The Gini coefficient for *Prdm1* cKO NP-specific Ig repertoire was significantly higher than that of WT cells (Fig. 3h) thereby providing evidence for clonal dominance. As a complementary Ig repertoire diversity metric, we computed Shannon entropy scores for the two distributions. In contrast with the Gini index, Shannon entropy increases with number of clonotypes and uniformity in their distributions. As expected, Shannon entropy values were lower for the *Prdm1* cKO repertoires (Fig. 3i). Together, these data show that, in the absence of Blimp-1, affinity maturation and clonal dominance are enhanced at the peak of the GC response (14 d.p.i.).

To test the increased cell seeding model for GC expansion caused by the loss of Blimp-1, we analyzed clonal diversity at 7 d.p.i., during the initiation of the GC response. If increased seeding was accounting for the phenotype, we expected to observe a broader initial BCR repertoire diversity in *Prdm1* cKO mice. Instead, we observed reduced clonal diversity at 7 d.p.i., indicating early clonal restriction (Extended Data Fig. 3a-c and Supplementary Table 7). This finding argues against increased GC seeding as a major mechanism underlying the *Prdm1* cKO phenotype.

### Blimp-1 constrains accessibility at ISRE and EICE motifs

To analyze the molecular consequences of Blimp-1 action on chromatin accessibility and to infer its target genes in GC states, we performed scMultiome-seq, jointly profiling gene expression and chromatin accessibility using NP^+^B220^+^ splenocytes isolated from WT and *Prdm1* cKO mice (14 d.p.i.). MIRA-based joint topic modeling^33^, which involves representation of each cell according to weighted mixtures of gene expression and chromatin accessibility revealed six discrete clusters, C1–C6 (Fig. 4a). Gene expression and chromatin accessibility topics that were specific for clusters C1-C6 are displayed (Fig. 4b,c) and their associated genes and chromatin coordinates are provided (Supplementary Table 8). Notably, a subset of C4 cells displayed the highest *Myc* gene signature score (Fig. 4d). To compare the scMultiome-defined clusters with those delineated by the scRNA-seq analysis, we calculated their ICGS2 gene signature scores, derived from the top 60 marker genes. Multiome clusters C1 and C2 exhibited the highest LZ-1 signature scores, C3 showed the highest LZ-2 scores, C4 showed the highest TZ scores, C5 exhibited the highest DZ-1 scores, and C6 exhibited the highest DZ-2 scores (Extended Data Fig. 4a-e). The corresponding violin plots demonstrate that scMultiome-derived clusters faithfully recapitulate the transcriptional signatures of the scRNA-seq–defined clusters in the WT cells. The scRNA-seq–defined clusters will be designated as “states” hereafter. Consistent with the cellHarmony analysis, which showed an increased relative representation of DZ over LZ GC B cells in *Prdm1* cKO mice, we observed altered proportions of scMultiome clusters based on ICGS2 signature scoring, further confirming that Blimp-1 deficiency skews GC composition toward the DZ fate (Extended Data Fig. 4f). Support vector machine (SVM) classification^34^ was used to more comprehensively analyze the correspondence between the six WT multiome clusters with the five reference scRNA-seq states (Fig. 4e-g and Extended Fig. 4g,h). Multiome clusters C1 and C2 were enriched for the LZ-1 state, C3 for LZ-2, C4 for TZ, C5 for DZ-1, and C6 for DZ-2. As was the case for scRNA-seq, multiome clusters derived from *Prdm1* cKO cells showed an overall correspondence with their WT counterparts, suggesting preservation of global GC state identities despite the perturbation. SVM classification was next used to infer alterations in relative frequencies of GC states because of the perturbation. The *Prdm1* cKO C3 cells manifested a reduction in relative frequency of LZ-2 classified cells with an increase in DZ-1 classified cells (Fig. 4h). Similarly, *Prdm1* cKO C4 cells displayed a reduction in relative frequency of TZ classified cells with an increase in DZ-1 cells (Fig. 4i). Thus, *Prdm1* cKO LZ-2 and TZ cells exhibited DZ-like features. Overall, these results were consistent with the scRNA-seq analysis, collectively demonstrating that Blimp-1 loss enhances LZ to TZ and TZ to DZ state transitions. Consistent with this interpretation, analysis of alterations in chromatin accessibility revealed a substantial increase in upregulated differentially accessible regions (upDARs) in the C3 and C4 *Prdm1* cKO clusters compared with C1 and C2 (Fig. 4j). Many of these upDARs were associated with G1 to S genes whose expression was increased in C3 and C4 *Prdm1* cKO cells (see below).

**Fig. 4.**
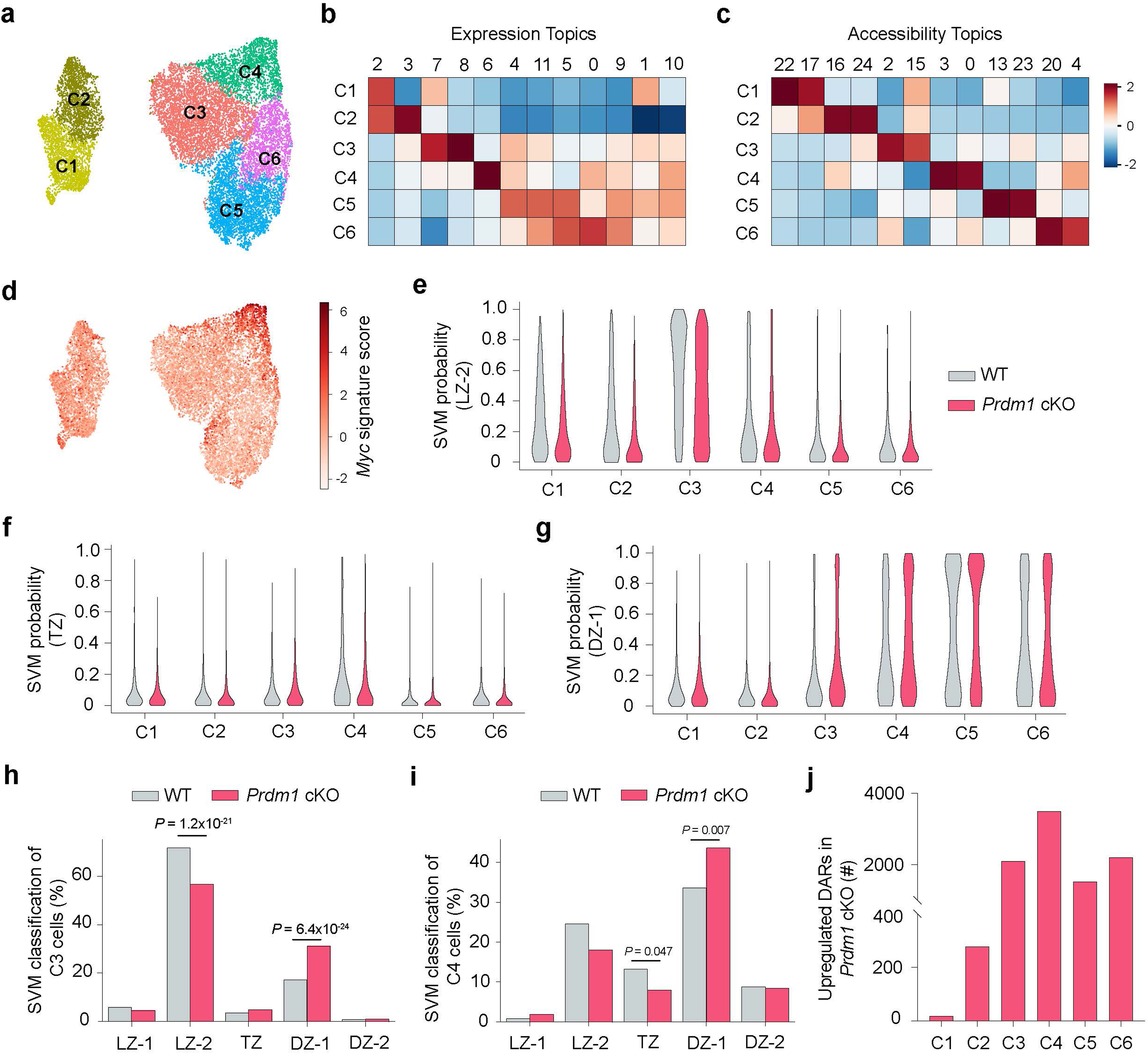
Multiome joint topic modelling reveals transcription and chromatin features underlying enhanced LZ-DZ transition in absence of Blimp-1. **a,** NP^+^B220^+^ cells isolated at 14 d.p.i. from WT and *Prdm1* cKO mice were profiled by scMultiome-seq (n=15,155). UMAP plot displaying 6 clusters (C1-6) was generated using MIRA and joint topic modelling of both RNA and ATAC features (WT=4,764 and *Prdm1* cKO=10,391). **b,c** Heatmaps showing relative prominence (z-score of average weight) of each gene expression **(b)** or accessibility **(c)** topic within designated cell clusters **(4a)**. **d,** UMAP displaying the *Myc* gene signature score across the multiome-seq clusters. **e-g,** Violin plots displaying the distribution of indicated ICGS2-delineated transcriptional states (y-axis) across the six scMultiome-seq defined clusters (x-axis) in WT and *Prdm1* cKO GC B cells. Probabilities were determined by SVM classification. **h,i** Bar plots displaying the frequency of WT and *Prdm1* cKO GC B cells within the C3 **(h)** and C4 **(i)** clusters that were assigned to indicated ICGS2-transcriptional states, as determined by SVM classification. **j,** Bar plot displaying the number of upregulated DARs for indicated scMultiome-seq clusters in *Prdm1* cKO compared with WT counterparts (14 d.p.i.). Statistical significance was tested by two-tailed Fisher’s Exact test 2×2 contingency table (**h,i**).

To analyze TF motif contributions to chromatin structure in WT GCBCs and their perturbations in the *Prdm1* cKO counterparts, at base-pair resolution, we utilized ChromBPNet^35^. The deep learning model predicts TF motif activity, while accounting for Tn5 transposase cutting bias. This approach enabled the systematic identification of TF motifs underlying chromatin accessibility changes during GCBC state transitions. The scMultiome dataset was used to train cluster-specific ChromBPNet models on pseudobulk ATAC-seq profiles derived from a given GCBC cluster. Each of these cluster-specific models was then used to generate corresponding base-pair resolution contribution score (CS) predictions across the union set of ATAC-seq peaks. These cluster-specific CS tracks revealed short DNA sequences, referred to as “seqlets”, that were predicted to be important in determining the accessibility profiles of their corresponding ATAC-seq peaks. Strikingly, many of the ChromBPNet seqlets corresponded to known TF motifs thereby implicating action of cognate TFs in binding to such sequences and controlling chromatin accessibility (Supplementary Table 9). Single nucleotide CS predictions for selected motifs, aggregated across all instances of their occurrence within accessible chromatin regions, are shown for each WT GCBC cluster (Fig. 5a). The analysis inferred dynamic action of multiple signaling induced TFs in regulating chromatin accessibility across GC B cell states with an increase in NFKB, MYC and POU activity preceding that of AP-1 (BATF).

**Fig. 5.**
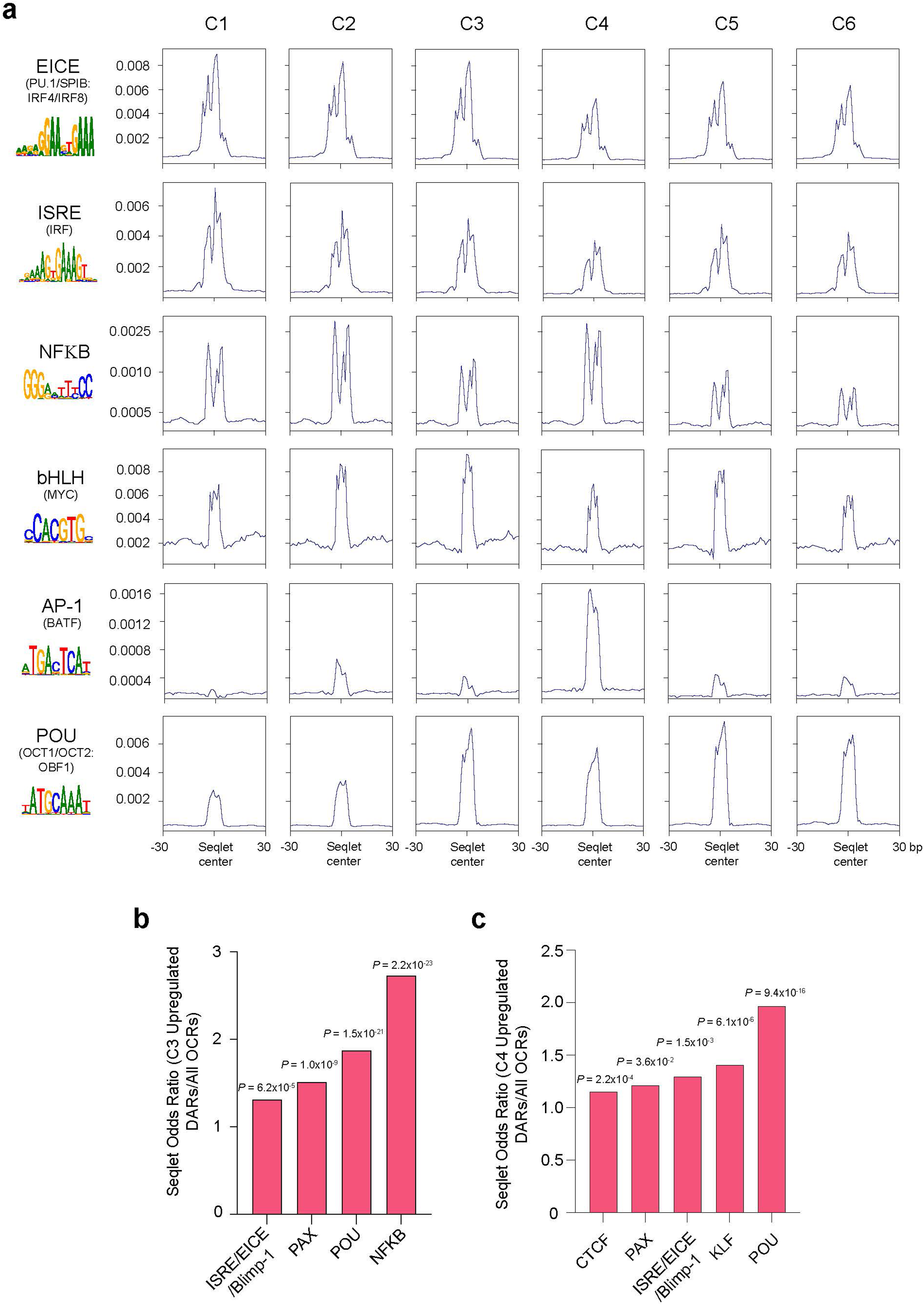
Loss of Blimp-1 enhances chromatin accessibility at PAX, POU and NFKB actionable regions during LZ to DZ transition. **a,** Aggregated ChromBPNet-derived CS signals at selected TF motifs for indicated WT GC B cell clusters within all peaks. Motif instances were identified by HOMER. Position weight matrix logos are displayed. **b,c** Bar plots displaying the relative increase in indicated seqlets predicted by ChromBPNet within the upregulated DARs in the C3 **(b)** and C4 **(c)** cluster of *Prdm1* cKO GC B cells. Statistical significance was tested by two-tailed Fisher’s Exact test and Benjamini-Hochberg method for correction of multiple-hypotheses testing (**b,c**).

We next measured the enrichment of TF motif family seqlets in the upregulated DARs of *Prdm1* cKO GC B cells (Supplementary Table 10). The upregulated C3 and C4 DARs were enriched for seqlets which matched EICE and ISRE motifs known to be bound by PU.1/SPIB:IRF4/IRF8 and IRF1/IRF2, respectively as well as Blimp-1 (Fig. 5b,c). Blimp-1 has been shown to antagonize the forementioned activators binding to EICE and ISRE motifs by recognition of an overlapping DNA sequence^36,37^. Additionally, we also observed enrichment for PAX, NFKB and POU motif seqlets in these regions, which predicted increased activities of their cognate TFs in the absence of Blimp-1. These findings suggested a chromatin gating function of Blimp-1 during the G1-S transition during which it antagonizes the action of constitutively expressed as well as signaling induced TFs in GC B cells.

### Blimp-1 represses expression of BCR-signaling genes in GC B cells

To distinguish the primary and secondary effects of loss of Blimp-1 on chromatin accessibility and gene expression, we sought to identify its direct transcriptional targets by integrating DEGs with ChromBPNet seqlet predictions and upDARs (Fig. 6a). A shared set of DEGs from the multiome-seq and scRNA-seq (Supplementary Table 11) enabled the identification of presumptive Blimp-1-repressed target genes. These genes (n=380) were upregulated in the absence of Blimp-1 and were associated with one or more ISRE, EICE or Blimp-1 motif seqlets within an OCR that was + 200 kb of their transcription start site (TSS) (Fig. 6b,c and Extended Data Fig. 5a-d). Within this target gene set we identified those that were associated with one or more upDARs (n=154) that in turn contained a Blimp-1-related motif seqlet (n=23) (Supplementary Table 11).

**Fig. 6.**
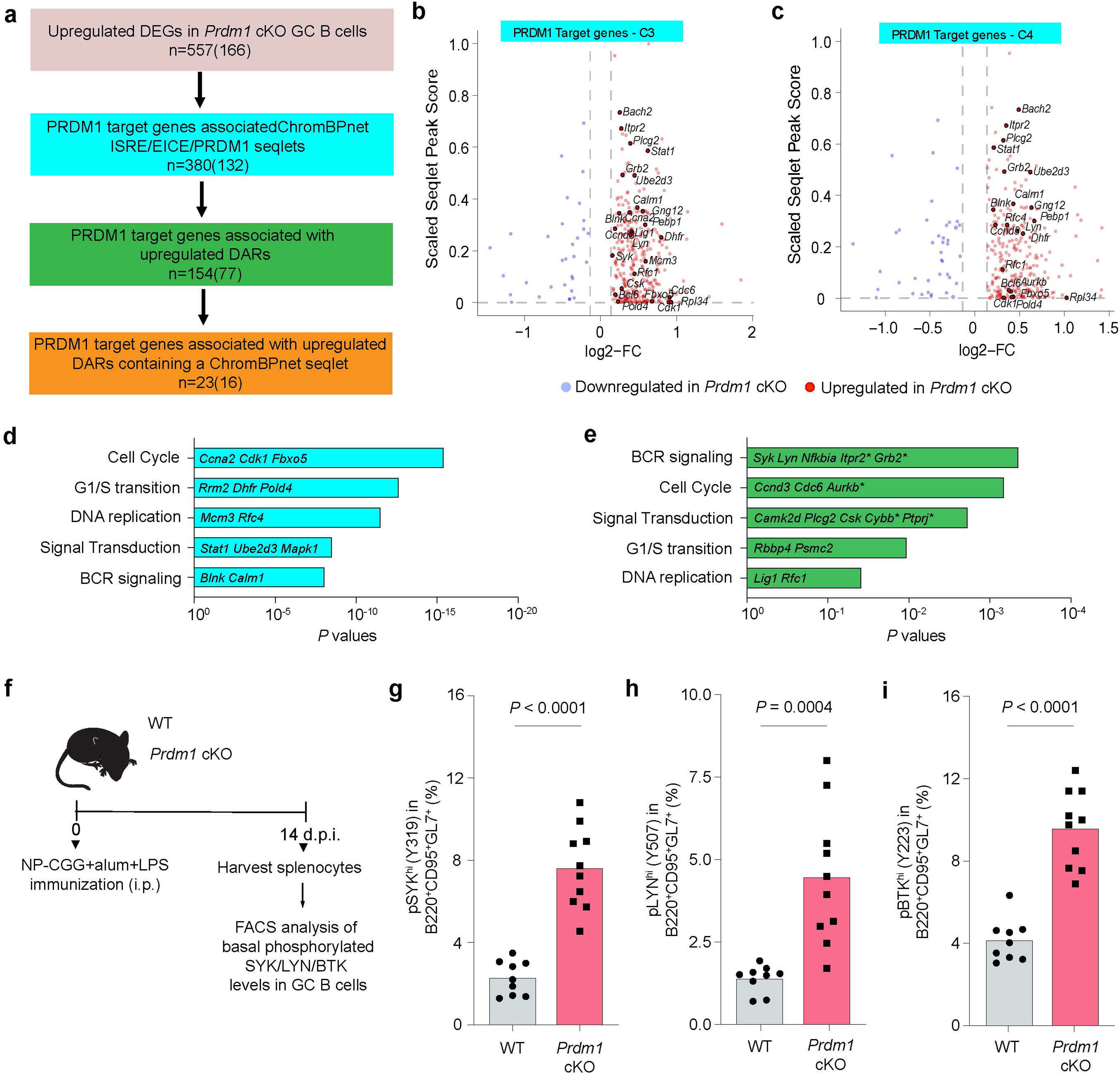
ChromBPNet modeling predicts Blimp-1 mediated repression of BCR signaling genes. **a,** Schematic rationale for identification of Blimp-1 target genes within upregulated DEGs in *Prdm1* cKO GC B cell clusters. Shared DEGS in scRNA-seq and multiome datasets (n=557) were used for downstream analysis. Color codes indicate target genes based on specified criteria. Blimp-1 target genes were associated (TSS +/-200kb) with one or more ISRE, EICE and Blimp-1 motif seqlets based on ChromBPNet models. These were annotated for association with one or more upDARs and ISRE, EICE and/or Blimp-1 motif seqlets located within them. Numbers outside parentheses denote the union of upregulated genes across the six clusters at each filtering step, whilst those in parentheses represent the genes overlaid with Blimp-1 ChIP-seq peaks from LPS activated B cells **(Methods)**. **b,c,** Volcano plots displaying the Blimp-1 target genes for the indicated clusters. Selected Blimp-1 target genes are displayed. **d,e,** Bar plots displaying key Reactome pathways associated with the Blimp-1 target genes identified by ChromBPNet seqlets **(d)** and ChromBPNet seqlets associated with one or more DARs **(e)**. Selected genes from the pathway enrichment analysis are highlighted. Asterisks indicate Blimp-1 target genes containing ISRE, EICE and/or Blimp-1 ChromBPNet seqlets within an associated upregulated DAR. **f,** Experimental design for phospho-flow analysis of pSYK (Y319), pLYN (Y507), and pBTK (Y223) in freshly isolated, unstimulated GC B cells from WT and *Prdm1* cKO mice (14 d.p.i.). **g-i,** Bar plots displaying the frequencies of pSYK^hi^ (Y319), pLYN^hi^ (Y507), and pBTK^hi^ (Y223) GC B cells from WT (n=9) and *Prdm1* cKO (n=10) mice (14 d.p.i.) respectively. Data in panels **g-i** are from two different experiments. Statistical significance was tested by one-tailed Fisher’s Exact test and Benjamini-Hochberg method for correction of multiple-hypotheses testing (**d,e**) and two-tailed Unpaired t-test (**g-i**).

To independently assess whether these targets could reflect direct Blimp-1 binding, we integrated our DEG and chromatin accessibility as well as ChromBPNet analyses with Blimp-1 ChIP-seq dataset from LPS activated B cells^36^. We recognized that the Blimp-1 binding regions were profiled in activated B cell/plasmablast contexts but reasoned that their convergence with our ChromBPNet analyses of Blimp-1 binding in accessible chromatin regions in GC B cells would nevertheless corroborate their delineation as direct targets. A substantial fraction of genes identified at each filtering step overlapped with Blimp-1 ChIP-seq peaks, including 16 of the 23 genes associated with Blimp-1-seqlet– containing upDARs (Fig. 6a and Supplementary Table 11, see Methods). Notably, these ChIP-validated targets included key components of the BCR signaling cascade e.g., *Syk, Lyn, Grb2, Itpr2*, and cell cycle regulators e.g., *Ccnd3, Mcm4, Rfc4, Pold4.* Blimp-1 binding was corroborated both at proximal and distal regulatory regions harboring ISRE, EICE, or Blimp-1 motifs. Gene set enrichment analysis of these Blimp-1 targets revealed significant overrepresentation of pathways involved in BCR signaling and G1 to S progression (Fig. 6d,e and Supplementary Table 12). The coordinated derepression of multiple genes within the BCR-signaling pathway, together with evidence for Blimp-1 occupancy at their regulatory elements, supports a model in which Blimp-1 directly constrains BCR signaling during GC selection.

To test whether Blimp-1-dependent repression of BCR signaling genes was reflected at the protein activity level, we performed phospho-flow cytometry at 14 d.p.i. We measured phosphorylated SYK (Y319), LYN (Y507), and BTK (Y223) in freshly isolated, unstimulated GC B cells. Subsets of *Prdm1* cKO GC B cells displayed significantly elevated baseline phosphorylation of all three kinases compared with WT cells (Fig. 6f-i and Extended Data Fig. 5e), demonstrating that the enhanced BCR-signaling potential predicted by chromatin accessibility changes was reflected by increased activities of multiple kinases downstream of the BCR. These results demonstrated that Blimp-1 functions to constrain BCR signaling.

## Discussion

As noted in the introduction, Conter et al. recently demonstrated a B-cell-intrinsic role for Blimp-1 in limiting GC expansion using bone marrow chimeras^22^. Our results strongly corroborate the following: increase in GC size, altered DZ:LZ ratios upon Blimp-1 loss and enhanced GC B cell proliferation. Conter et al. observed ∼2-fold elevation in MYC protein and increased S-phase fractions, aligning with our finding of elevated MYC target signatures and precocious G1-S programming in *Prdm1* cKO cells. The absence of increased *Myc* transcripts in both datasets suggests post-transcriptional or post-translational impact on MYC protein expression caused by the Blimp-1 perturbation. Our work significantly extends the shared findings through the integrated scRNA-seq, scMultiome and BCR-seq analyses as follows: (1) demonstrating increased clonal dominance and reduced BCR repertoire diversity by analyzing Gini indices and Shannon entropy (2) revealing chromatin-level mechanisms through which Blimp-1 gates accessibility at ISRE, EICE, NF-κB, and POU motifs at regulatory regions associated with BCR signaling and cell cycle genes; (3) corroboration of Blimp-1 target genes via ChIP-seq integration (4) evidence for enhanced BCR signaling in Blimp-1 lacking GC B cells through ex vivo phospho-flow and (5) independent functioning of Blimp-1 and NR4A1 GC feedback pathways. Thus, we establish Blimp-1 as a central feedback regulator that constrains GC amplitude and shapes clonal diversity through chromatin gating of BCR signaling.

Our findings on clonal diversity demonstrate that at 7 d.p.i., despite elevated usage of the NP-biased IGHV1-72 gene segment, *Prdm1* cKO mice exhibit reduced clonal diversity. This early restriction indicates that the exaggerated GC phenotype arises from preferential expansion of a limited NP⁺ repertoire, rather than increased recruitment of diverse clones. If GC seeding were enhanced, the initial repertoire would be expected to broaden, resulting in increased, not reduced, diversity in the absence of Blimp-1. The progression from restricted diversity at 7 d.p.i. to pronounced clonal dominance at 14 d.p.i. therefore strongly supports enhanced clonal amplification within the GC over increased seeding. Although we cannot exclude a partial contribution from attenuated GC exit, see below, our data and prior studies^32,38^ indicate that only a small fraction of GC B cells are poised to differentiate into PC precursors during the peak of the NP response (14 d.p.i.). Thus, GC B cell expansion in *Prdm1* cKO mice at this stage is unlikely to be due to their inability to exit the GC. While DZ to LZ migration is indeed more frequent than LZ to DZ recycling^39^, even modest enhancement of the LZ to DZ transition rate can substantially impact GC dynamics when compounded over multiple rounds. Our model proposes Blimp-1 constrains this specific transition through signaling attenuation, which is conceptually distinct from blocking the PC exit pathway.

In agreement with the model proposed by Ise et al.^31^, in which the IRF4:BCL6 ratio determines PC fating versus recycling in the GC, we observed decreased *Irf4* and *Cd69* transcripts with elevated *Bcl6* expression (a Blimp-1 target). Our results are consistent with diminished frequency of IRF4^hi^BCL6^lo^ GC B cells in *Prdm1* cKO mice reported by Conter et al.^22^. Importantly, Conter et al., demonstrated that Blimp-1 is required to downregulate BCL6 and induce IRF4 in GC B cells in response to IL-21 and CD40 signaling *ex vivo*. Conversely, BCL6 directly represses *Prdm1*, forming a reciprocal antagonistic circuit that enforces GC fate and delays PC differentiation^40^. Thus, Blimp-1 loss can generate a dual effect, enhanced BCR signaling promoting stronger positive selection, while the reduced IRF4:BCL6 ratio favors GC retention rather than exit. Impaired GC exit may play a more prominent role later in the GC response, as PC precursors begin to accumulate.

Our analysis has revealed an unsuspected function of Blimp-1 as a feedback attenuator of the GC B cell response. We propose that transient and sub threshold levels of Blimp-1, that are induced by BCR and CD40 signaling, feedback to inhibit signal transduction through these pathways and in so doing lower clonal dominance that can accompany affinity maturation. ChromBPNet analysis of chromatin accessibility in *Prdm1* cKO GC B cells delineated candidate upregulated target genes within BCR-signaling pathways e.g., *Syk, Lyn, Grb2, Itpr2* whose regulatory regions harbor ISRE/EICE/Blimp-1 seqlets contributing to open chromatin. We propose that Blimp-1 restrains transcription at these elements in part by opposing IRF and ETS family activators e.g., PU.1/SPIB:IRF8^36,37^. Enhanced BCR signaling via modest yet coordinated increases in expression of multiple genes encoding signaling components would be expected to elevate signaling-induced TFs. In line with such a signaling-amplification model, upDARs in *Prdm1* cKO LZ cells were enriched for NF-κB and POU (Oct) seqlets, consistent with increased activity of the cognate TFs. Recent work indicates that the NF-κB p52–ETS1 complex promotes GC formation by inducing POU2F1 (OCT1) and POU2AF1 (OBF1)^17^, factors that are required to sustain BCR signaling and GC maintenance^15^. We identified *Pou2af1* as a presumptive Blimp-1 target gene, it displayed increased expression in *Prdm1*-deficient GC B cells, suggesting that Blimp-1 restrains POU activity in part via direct repression of *Pou2af1*. Notably, *Prdm1* expression is reduced in p52–ETS1-deficient GC B cells, consistent with a model in which BCR/CD40-driven NF-κB activity transiently induces Blimp-1 as well as *Pou2af1*. Taken together, the p52–ETS-POU (positive) and p52–ETS-Blimp-1 (negative) gene regulatory interactions form an incoherent module that tunes GC amplitude and diversity, reminiscent of the previously proposed IRF4–BCL6–Blimp-1 incoherent feed-forward architecture^41^. How these two regulatory modules interface with *Nr4a1* remains to be more fully explored. *Nr4a1* is an immediate early gene that is induced by BCR signaling^18^ whereas *Prdm1* appears to manifest slower signaling dependent induction^21^. Notably, we did not observe diminished expression of *Nr4a1* in *Prdm1*-deficient GC B cells indicating that Blimp-1 action in tuning GC amplitude and Ig repertoire diversification is not mediated through controlling the expression of *Nr4a1*.

The coordinate expression of *Nr4a1* and *Prdm1* in activated LZ-1 cells, despite their independent regulation, suggests parallel negative feedback pathways that together restrain GC amplitude. *Nr4a1* is an immediate early gene induced rapidly upon BCR signaling, whereas *Prdm1* exhibits slower, sustained induction. Their co-expression may reflect hierarchical temporal control: NR4A1 providing immediate feedback to prevent hyperactivation, while Blimp-1 enforces a more durable chromatin-level constraint. The observation that *Nr4a1* expression is unaffected in *Prdm1* cKO cells and vice versa indicates these pathways function independently, raising the question of whether dual targeting might synergistically modulate GC responses. How these parallel feedback systems are coordinated and whether they gate distinct aspects of the selection response, represents an important area for future investigation.

Strong Tfh–B-cell interactions promote PC differentiation via *Prdm1* induction, whereas weaker interactions favor recycling^31^; our data extend this paradigm by showing that Blimp-1 also restricts GC expansion by gating the LZ to DZ transition. Our multiome analyses point to Blimp-1-dependent chromatin gating as a direct means to tune expression of BCR-signaling genes; modest, coordinated upregulation of components such as *Syk*, *Lyn*, and *Btk* in the absence of Blimp-1 likely contributes to their increased activities resulting in augmented NF-κB and POU chromatin action, promoting G1–S entry and DZ proliferation. We propose that transient, NF-κB-linked Blimp-1 induction in the LZ feeds back to dampen signaling components and constrain NF-κB/POU activity, thereby limiting LZ to DZ flux. In the absence of Blimp-1, positively selected clones undergo premature G1–S commitment and increased DZ proliferation. Thus, Blimp-1 functions as a molecular brake on GC progression that limits clonal dominance and shapes repertoire diversity.

Our integrative analyses indicate that Blimp-1 regulates a broad network of targets in parallel, including components of BCR signaling, cell cycle progression, and transcriptional regulation. Given this coordinated regulation, perturbation of any single downstream gene would be unlikely to rescue the full phenotype, and failure to observe rescue would be difficult to interpret. Meaningful genetic rescue would likely require combinatorial or dosage-controlled perturbations. While our *Cd19*-Cre model establishes a B-cell–intrinsic role for Blimp-1 in restraining GC dynamics, inducible or activation-stage–restricted deletion will be essential to dissect when and where Blimp-1 acts within the GC, and to determine whether its dominant effects occur in the LZ, TZ, or during GC exit.

Our findings have implications for modulating GC responses in health and disease. By revealing that Blimp-1 constrains BCR signaling strength, regulates LZ to DZ cycling, and limits clonal amplification, our study identifies regulatory nodes that could be leveraged to shape humoral immunity. Temporally tuning Blimp-1-dependent pathways may enhance vaccine efficacy for pathogens that require sustained GC engagement and broad epitope coverage. For example, transient modulation of these molecular pathways during prime-boost regimens could prolong GC persistence while maintaining repertoire diversity, potentially improving responses to HIV or influenza vaccines where both affinity and breadth are critical. Conversely, the pathways derepressed by the loss of Blimp-1 (BCR signaling components, NF-κB/POU/IRF activity) represent intervention points for conditions where excessive GC activity drives pathology. In systemic lupus erythematosus, where BCL6 is constitutively active, enhancing Blimp-1 function or targeting its downstream effectors (*Syk*, *Lyn*, *Pou2af1*) could dampen pathogenic GC responses. Small-molecule modulators of the chromatin-encoded checkpoint identified here, particularly those affecting ISRE/EICE accessibility or NF-κB/POU activity, could provide tunable control over GC magnitude and diversity.

## Acknowledgments

We thank the expert personnel within Division of Laboratory Animal Resources, the Flow Cytometry Core, the Single Cell Core and UPMC Genome Center at the University of Pittsburgh for their invaluable assistance. We particularly wish to acknowledge the Lafyatis lab for help with the scRNA-seq and BCR-seq library preparations and Stuart Hay for setting up the viewer for scRNA-seq datasets. This research was supported by the UPMC ITTC fund and NIH grants (RO1 AI145064; H.S.) and (RO1 AI114587; D.M.R.).

## Author contributions

H.S. and D.M.R. conceived and supervised the study. G.K.M.V. and B.R. performed all experiments. G.K.M.V., S.G., N.P., and P.H.G. performed the computational analysis. D.C., K.T., and S.K. helped with select computational analyses. N.P. and L.M.H. helped with the experiments. H.X., N.S., J.D. and D.M.R. provided critical inputs during the study. G.K.M.V. and H.S. wrote the manuscript with input from other authors.

## Tables

NI-RS42391_Table-1

NI-RS42391_Table-2

NI-RS42391_Table-3

NI-RS42391_Table-4

NI-RS42391_Table-5

NI-RS42391_Table-6

NI-RS42391_Table-7

NI-RS42391_Table-8

NI-RS42391_Table-9

NI-RS42391_Table-10

NI-RS42391_Table-11

NI-RS42391_Table-12

NI-RS42391_Table-13

## Methods

### Mice

*Prdm1*^fl/fl^ (008100) and *Cd19*^cre/cre^ (006785) mice were purchased from Jackson laboratory and were intercrossed to generate *Prdm1*^fl/fl^*Cd19*^cre/+^ (*Prdm1* cKO) and *Prdm1*^+/+^ *Cd19*^cre/+^ (WT) littermates. Mice were housed under specific pathogen-free conditions (20– 26 °C; 30–70% relative humidity; 12-h light/dark cycle). All procedures complied with University of Pittsburgh IACUC protocols 21059197 and 24054803.

### Immunization

Eight to ten week old *Prdm1* cKO and WT mice were immunized intraperitoneally with 50 μg NP(23)-CGG (Biosearch Technologies) mixed with 50% (v/v) imject alum (Thermo Fisher Scientific) and 1 μg LPS (Sigma). All experiments used both male and female mice, with sexes balanced across genotypes and experimental cohorts. We did not observe sex-specific differences in GC magnitude (cell numbers or frequencies), transcriptional profiles, or clonal features. No formal randomization was used to assign animals or samples to experimental groups. Animals were allocated according to genotype and experimental availability.

### Quality control of intraperitoneal immunizations

To ensure the success and consistency of intraperitoneal immunizations, we implemented a quality-control strategy based on GC responses (see below for staining methods). Since *Prdm1* cKO mice fail to mount detectable serum antibody responses, NP-specific antibody titers could not be used as a reliable indicator of immunization efficacy. Instead, GC induction was used as a functional readout of successful antigen delivery and immune activation. Baseline GC B cell numbers were first established in unimmunized WT and *Prdm1* cKO mice analyzed in parallel. For each genotype, the mean GC B cell number under steady-state conditions was calculated and used as a minimum threshold. Immunized mice whose GC B cell numbers fell at or below this genotype-matched baseline were prospectively designated for exclusion, as such values would indicate a failure of effective immunization. All immunized mice included in the study exhibited robust GC responses that exceeded the predefined baseline threshold, and therefore no animals were excluded on this basis. This quality-control criterion was applied uniformly across all experiments and time points.

### Flow cytometry

Spleens were dissociated to single-cell suspensions in PBS (Ca²⁺/Mg²⁺-free) + 2% FBS + 2 mM EDTA (FACS buffer). Erythrocytes were lysed in ACK lysis buffer (Thermo Fisher) for 5 min at 23°C, quenched in FACS buffer, and washed. After a brief PBS wash to remove serum proteins, cells were stained with Zombie NIR (BioLegend) and anti-CD16/32 (2.4G2, 25 µg/mL) for 10 min on ice, followed by fluorochrome-conjugated antibodies (listed in Supplementary Table 13) for 20 min on ice. Standard doublet exclusion (FSC-A/H, SSC-A/H), FMO/compensation controls, and NP binding controls (unconjugated CGG; NP-negative samples) were used. Cells were fixed in BD Cytofix (#554655) for 20 min at 4 °C, washed, and resuspended in PBS. Data were acquired on a Cytek Aurora and analyzed in FlowJo (Tree Star). GC B cells were gated as B220⁺ GL-7⁺ Fas⁺, with NP⁺ subsets as indicated.

### Phospho-flow staining

Single-cell suspensions were prepared from spleens in pre-warmed B cell medium (RPMI 1640 supplemented with 10% fetal bovine serum, GlutaMAX or L-glutamine, penicillin–streptomycin, HEPES (pH7.2), and 50 µM 2-mercaptoethanol) and resuspended at 5 × 10⁶ cells ml⁻¹ in 1 ml volumes in round-bottom tubes. Cells were washed once with PBS before resuspension and were rested at 37 °C for 15 minutes. For fixation and permeabilization, one volume of 2× Cytofix/Perm/Wash buffer (prepared by diluting paraformaldehyde and BD Perm/Wash buffer) was added directly to unstimulated cells, followed by incubation at 23°C for 15 minutes and on ice for 30–60 minutes. Cells were washed twice with 1× Perm/Wash buffer. Cells were incubated with Fc receptor–blocking antibody (anti-CD16/CD32, clone 2.4G2; 1:200) for 5 min at 4 °C, followed by surface and/or intracellular phospho-specific antibody staining (listed in Supplementary Table 13) in Perm/Wash buffer for 60 minutes at 4 °C in the dark. After final washes, cells were resuspended in staining buffer and analyzed by flow cytometry.

### Isolation of NP^+^B220^+^ cells

For sorting, splenocytes (7 and 14 d.p.i.) were stained with eFluor780 live/dead, B220 (RA3-6B2; PE-Dazzle 594), and NP–PE (NP(30)–PE) for 20 min at 4 °C and sorted as NP⁺B220⁺ on a FACSAria II (70 µm nozzle; 4 °C). Post-sort purity were above 95%.

### scRNA-seq and BCR-seq

Sorted NP⁺B220⁺ cells were suspended in 5% FBS and loaded on a Chromium Controller (10x Genomics; Single Cell 5′ v2). Samples were collected from WT (n=3) and *Prdm1* cKO (n=3) mice at 14 d.p.i. and WT (n=1) and *Prdm1* cKO (n=1) mice at 7 d.p.i. Libraries were prepared as per the manufacturer’s protocol as described previously^32^. From the cDNA, one aliquot was used for gene-expression library prep; a second was amplified for VDJ using the BCR kit. Libraries were sequenced on an Illumina NovaSeq 6000 with manufacturer recommended read structures for 5′ GEX and VDJ. Target depths were ∼30k reads/cell (GEX) and ∼5k reads/cell (VDJ).

### scRNA-seq data processing

FASTQs were processed with Cell Ranger v6.1.2 using the mm10 reference. UMI count matrices from WT 14 d.p.i. NP⁺B220⁺ cells (HDF5) were analyzed with AltAnalyze ICGS2 (default parameters) to delineate clusters and marker genes^25^. Initial cell state labels were assigned by ICGS2/GO-Elite enrichment and refined by literature markers. Non-B-lineage contaminants were removed.

### scRNA-Seq annotation and integration

WT clusters served as reference states for supervised alignment of *Prdm1* cKO cells using cellHarmony^30^ (centroid-based label transfer; Pearson r ≥ 0.7 on log₂-scaled counts). Fisher’s exact test was performed to assess differential cell type frequency.

### Pseudotime trajectory and RNA velocity analyses

Cell-by-gene counts and cell type labels file were filtered for cells from LZ-1, LZ-2, TZ, DZ-1, and DZ-2 clusters, and were used as input for Monocle2^29^ pseudotime analysis using default options. LZ-1 GC B cells were designated as the root for the analysis. The counts data was modeled with negative binomial distribution using Monocle 2 (‘expressionFamily=negbinomial.size’). Monocle 2 was allowed to select its genes for pseudotime estimation based on differential gene analysis across the filtered B-lineage cells (‘fullModelFormulaStr = ∼Groups’). The reverse graph embedding (RGE) (‘method’ in reduceDimension) method was set to “DDRTree” as recommended. The Monocle2 cell state that contained the maximum number of LZ-1 GC B cells was set as the “root” state to calculate the pseudotime of all cells. Velocyto v0.17.17^28^ was used to generate loom files with spliced/unspliced counts for LZ, TZ, and DZ clusters. Pre-processing and dynamical modeling used scVelo v0.3.3^42^. Scanpy v1.11.4^43^ with default settings was used to generate velocity streams over the UMAP manifold.

### Gene Expression Analyses

Expression values for *Prdm1* and *Nr4a1* were normalized to counts-per-10k and log1p-transformed “expressing” cells were defined from the counts layer as >0. Analyses were stratified by GC states (LZ-1, LZ-2, TZ, DZ-1, DZ-2) and visualized with violin plots.

### Gene signature score

To calculate gene signature scores, an algorithm from our previous study^7^ was adopted to control the variation of quality and complexity of scRNA-seq data of individual cells. To calculate the ICGS2 gene signature scores, this algorithm was modified based on the top 60 marker genes for each of the scRNA-seq clusters.

### Differential gene expression analyses

For scRNA-seq datasets, DEG analysis was performed using cellHarmony between the reference (WT) and the query (*Prdm1* cKO) datasets. Genes with an absolute fold > 1.1 (log₂ fold change ≥ 0.13) and empirical Bayes t-test p-value < 0.05 (Benjamini Hochberg corrected) were considered to be differentially expressed. For multiome-seq data, differential gene expression analysis was performed using Scanpy (v1.10.3) between reference (WT) and query (*Prdm1* cKO) cells for each cluster. Genes with a Benjamini-Hochberg correct pooled-variance Student’s t-test P value < 0.05 and absolute log₂ fold change ≥ 0.13 were considered to be differentially expressed.

### Pathway enrichment analysis

Pathway enrichment analysis was performed using Enrichr (Ma’ayan Laboratory; https://maayanlab.cloud/Enrichr/). The top 60 marker genes identified for each indicated cluster and DEGs were analyzed using Enrichr. Functional enrichment was queried against the Reactome and Gene Ontology (GO) Biological Process libraries. TF–associated enrichment was assessed using the ChEA library. Enrichr calculates enrichment using Fisher’s exact test and reports nominal *P* values and Benjamini– Hochberg–adjusted *P* values; adjusted *P* values were used to determine statistical significance.

### BCR analysis of 5’-end scRNA-seq

VDJ FASTQs were processed with Cell Ranger VDJ v6.1.2 with GRCm38 mouse reference. BCR and GEX barcodes were linked to couple clonotypes with transcriptomes. NP-specific clones were defined as cells bearing IGHV1-72*01 (VH186.2) rearrangements; cells lacking paired GEX–VDJ were excluded. Clonotypes were defined from the top-UMI productive IGH and IGK/IGL chains using CDR3 amino-acid sequence + V + J genes (alleles collapsed), as described¹,⁷. SHM rates and W33L calls were as described^7^.

### BCR Clonotype Analysis

Within each sample, all clonotypes were enumerated based on their paired IgH and IgL sequences and then cells were grouped by clonotype. Lorenz curves plotted cumulative cells versus cumulative clonotypes. To equalize depth for estimation of Gini coefficients and Shannon entropy scores, clone sizes were rarefied to the minimum paired-cell count across samples (subsampling without replacement). For each replicate, the Gini coefficient^44^ and Shannon entropy scores^45^ were computed from clone counts and summarized per-sample rarefied means with 95% CIs (2.5–97.5th percentiles of replicate distributions). Per-genotype values are the mean of experiment-level rarefied Gini indices and Shannon entropy scores, weighted equally across experiments.

### scMultiome-seq

Sorted NP^+^B220^+^ splenocytes were washed twice with PBS containing 0.04% BSA and lysed in 100 μL of chilled lysis buffer [10 mM Tris-HCl (pH7.4), 10 mM NaCl, 3 mM MgCl₂, 0.1% Tween-20, 0.1% IGEPAL, 0.01% digitonin, 1% BSA, 1 mM DTT, 1 U/μL RNase inhibitor (Protector, Sigma)]. Cells were mixed by pipetting and incubated on ice for 3 min. Lysis was quenched with 1 mL of chilled wash buffer (same composition as lysis buffer excluding digitonin) and mixed gently. Nuclei were washed three times with wash buffer and resuspended in diluted nuclei buffer (10x Genomics) supplemented with 1 mM DTT and 1 U/μL RNase inhibitor. Nuclei were counted, resuspended in chilled PBS (750–1,000 nuclei/μL), and aliquoted (180 μL per sample). Labeling reactions were quenched with 2% BSA in PBS, followed by centrifugation at 500 × g for 4 min at 4 °C. After pooling and washing, nuclei were resuspended in diluted nuclei buffer and processed according to the Chromium Next GEM Single Cell Multiome ATAC + Gene Expression User Guide (10x Genomics) through the pre-amplification PCR step. Ten microliters of the pre-amplified product were fractionated using 0.6× SPRI bead cleanup to separate cDNA (gene expression) and gDNA (ATAC) products. Final libraries were prepared according to the manufacturer’s protocol and sequenced on a NovaSeq 6000 (Illumina) obtaining a depth of 30,000 reads for the mRNA and ATAC libraries.

### scMultiome data analysis

FASTQ files were processed with Cell Ranger ARC (v2.0.2) using the GRCm38 (2011) reference genome for alignment. Outputs from individual GEM wells were aggregated using the ‘aggr’ function. Cell and gene filtering were performed in Scanpy (1.11.4), retaining cells with >2000 detected genes and <10% mitochondrial reads, and genes expressed in at least 20 cells. Cell cycle effects were regressed out using the ‘regress_out’ function in SCANPY with G1/S and G2/M gene sets. MIRA v2.1.1 was used for joint topic modeling of expression and accessibility. A joint embedding integrated topic weights, followed by UMAP and Leiden clustering. Peaks were called with MACS2 v3.0.3 (see below).

### SVM based cell state predictions

A multiclass SVM (scikit-learn v1.3.2) was trained on standardized gene-expression features from the scRNA-seq reference states (ICGS2 labels) and applied to the Multiome dataset to infer LZ-1, LZ-2, TZ, DZ-1, DZ-2 probabilities per cell. The maximum-probability class was assigned as the label; probability distributions were retained for statistics^34^. Receiver operating characteristic curves (using Youden’s J statistic calculated per Leiden cluster) defined high-confidence probability cutoffs for each state.

### MIRA Topic Modeling

MIRA (v2.1.1)^33^ topic modeling was applied to both RNA expression and ATAC accessibility data, inferring topic weights for each cell. Weights were averaged per cluster to build a cluster × topic matrix, then each topic was z-score normalized across clusters. The topics showing the highest variance between clusters were selected as cluster specific topics.

### ChromBPNet Models and Analyses

Cluster/genotype-specific pseudo-bulk BAMs were generated from single-cell multiome dataset by using Sinto’s ‘filterbarcodes’ function and used to call peaks with MACS2 (v2.2.7.1). These peak sets were merged and used to train cluster/genotype-specific ChromBPNet (v0.1.7) models^35^. A Tn5 bias model was trained by running ChromBPNet ‘bias pipeline’ with nonpeak sequences from inaccessible chromatin regions. Bias-corrected BigWig files were generated by running ChromBPNet ‘pred_bw’ and base-pair resolution contribution score predictions were generated by running ChromBPNet ‘contribs_bw’ with the top accessibility topics (MIRA) per cluster, and cluster specific upregulated DAR peak sets. The resulting profile scores AnnData objects were used as input into TFModisco ‘motifs’ function using sampling of 10,000 seqlets and the ‘report’ function using the Jaspar 2024 Core Vertebrates (non-redundant) PFM collection. Seqlets matching motifs in the same Jaspar motif cluster family were merged for quantification of TF motif family seqlet instances sampled from either the union set of peaks or cluster specific upregulated DARs.

### Differential Chromatin Accessibility Analyses

scATAC data (multiome) were normalized and log transformed using Scanpy (v1.10.3)^43^. Peaks were annotated with genomic feature information. Differential chromatin accessibility was calculated using Muon v0.1.6 between *Prdm1* cKO and WT cells within each Leiden cluster. Peaks with a false discovery rate < 0.05 and absolute log₂ fold change ≥ 0.13 were classified as differentially accessible regions (DARs)^46^.

### Blimp-1 Candidate Target Gene Analysis

HOMER annotatePeaks.pl was used to identify ATAC peaks harboring ISRE/EICE/Blimp-1 motifs. For each motif instance, the dot product between the ChromBPNet contribution-score vector and the motif’s position-weight matrix was computed; instances with positive mean dot product were classified as seqlet hits. Genomic coordinates of seqlet hits were intersected with ATAC peaks linked to cluster DEG lists. Two additional notations were added to candidate Blimp-1-repressed genes: (i) overlap with ≥1 up-DAR with the gene body; and (ii) co-localization of a Blimp-1-related seqlet and an up-DAR.

### Integration with Blimp-1 ChIP–seq datasets

Published Blimp-1 ChIP–seq data from LPS activated B cells/plasmablasts were used to assess Blimp-1 occupancy at candidate target genes identified in this study^36^. Blimp-1 ChIP–seq peak coordinates were intersected with gene sets from peak-gene linkage analysis using 10x peak annotations data and MACS2 peak calling from the ChromBPNet pipeline to determine overlap with Blimp-1-bound regions. Intersections were performed at successive levels of stringency corresponding to the filtering steps applied in the RNA and chromatin analyses, including genes upregulated in *Prdm1* cKO GC B cells, genes associated with ChromBPNet predicted ISRE-, EICE-, or Blimp-1-seqlets, and genes associated with upregulated DARs in *Prdm1* cKO GC B cells. Genes overlapping Blimp-1 ChIP–seq peaks at each step were retained as Blimp-1-bound candidates.

### Statistical analysis

Data collection and analysis were not performed blind to the conditions of the experiments. No statistical methods were used to pre-determine sample sizes but our sample sizes are similar to those reported in previous publications^7,32^. Data distribution was formally assessed for normality prior to statistical analysis. Homogeneity of variances was evaluated where applicable. Based on these assessments, either parametric or nonparametric tests were applied as appropriate. All statistical tests were two-tailed unless otherwise specified. Pathway enrichment analyses performed using Enrichr were conducted using the statistical framework implemented in the Enrichr platform. For the biological datasets, we used Prism version 10 (GraphPad) for differential testing. Pairwise statistical tests were performed using a Student’s t-test or Mann-Whitney test, whereas multicondition comparisons were performed using a one-way ANOVA with Tukey’s multiple comparison test or Kruskal Wallis with Dunn’s multiple comparison test based on the distribution, as indicated. Difference in proportions were calculated using Fisher exact test and Benjamini-Hochberg method for correction for multiple hypotheses testing. Two-sample Kolmogorov– Smirnov tests were used to compare the state-probability distributions between *Prdm1* cKO and WT cells within each Leiden cluster (significance at *P* < 0.05). Aggregate motif counts in upDARs versus background peaks from multiome-seq were tabulated per cluster. Fisher’s exact tests (SciPy v1.9) with Benjamini–Hochberg correction were used for identifying enriched motifs. Co-expression in double-positive cells was assessed with Spearman correlations (95% CIs via Fisher and a cluster-blocked permutation p-value). Co-detection enrichment used Fisher’s exact on 2×2 tables, and detection prevalence by state was summarized with Wilson 95% CIs plus global chi-square/trend tests and pairwise Fisher (odds ratios).

## Data availability

All raw and processed data associated with the scRNA-seq analyses have been deposited in GEO: GSE305877. Flow cytometry data underlying the figures are provided as source data.

## Code availability

The R code defining the clones and all other codes are provided at (https://github.com/pitt-csi).

---

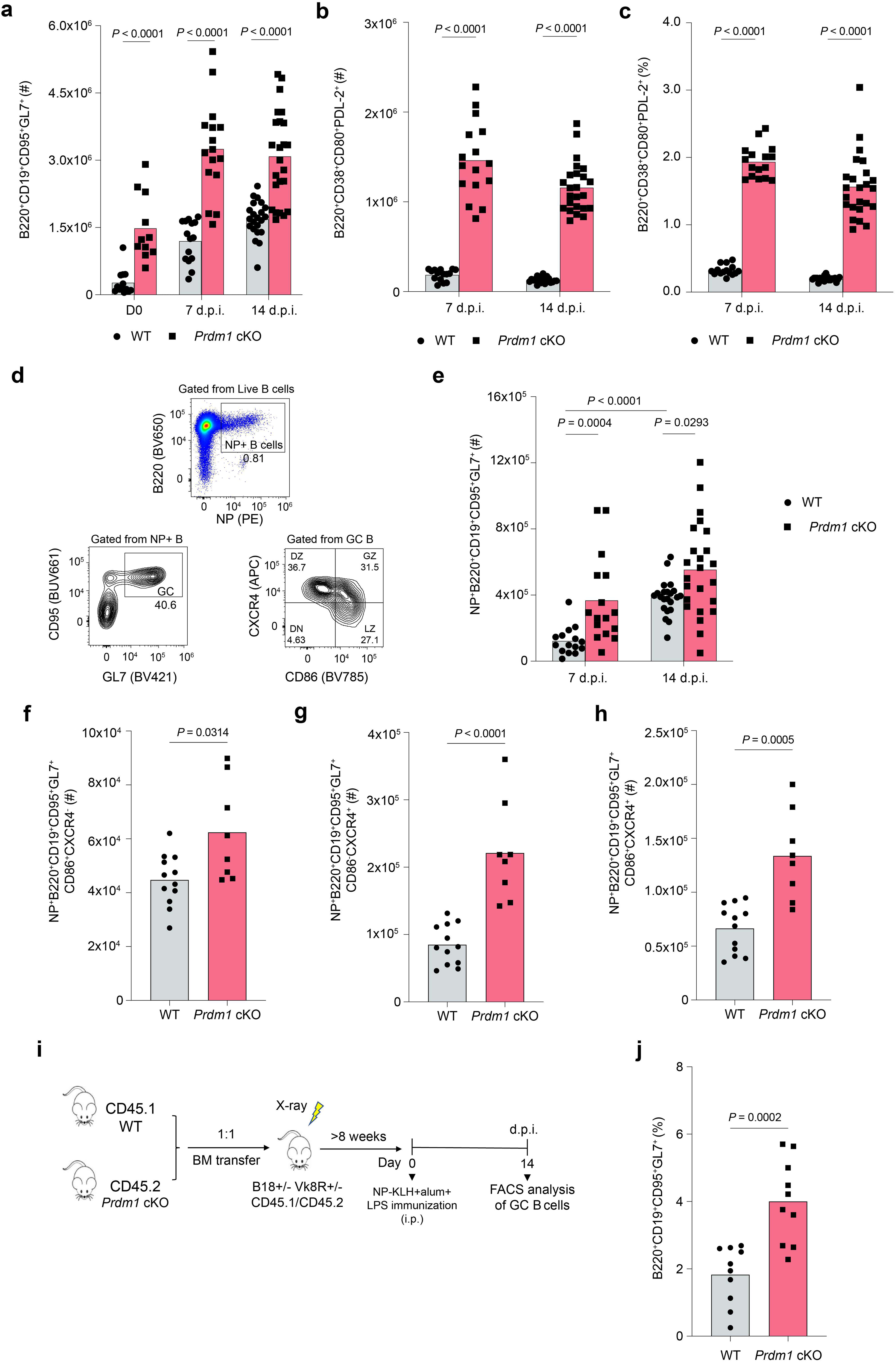

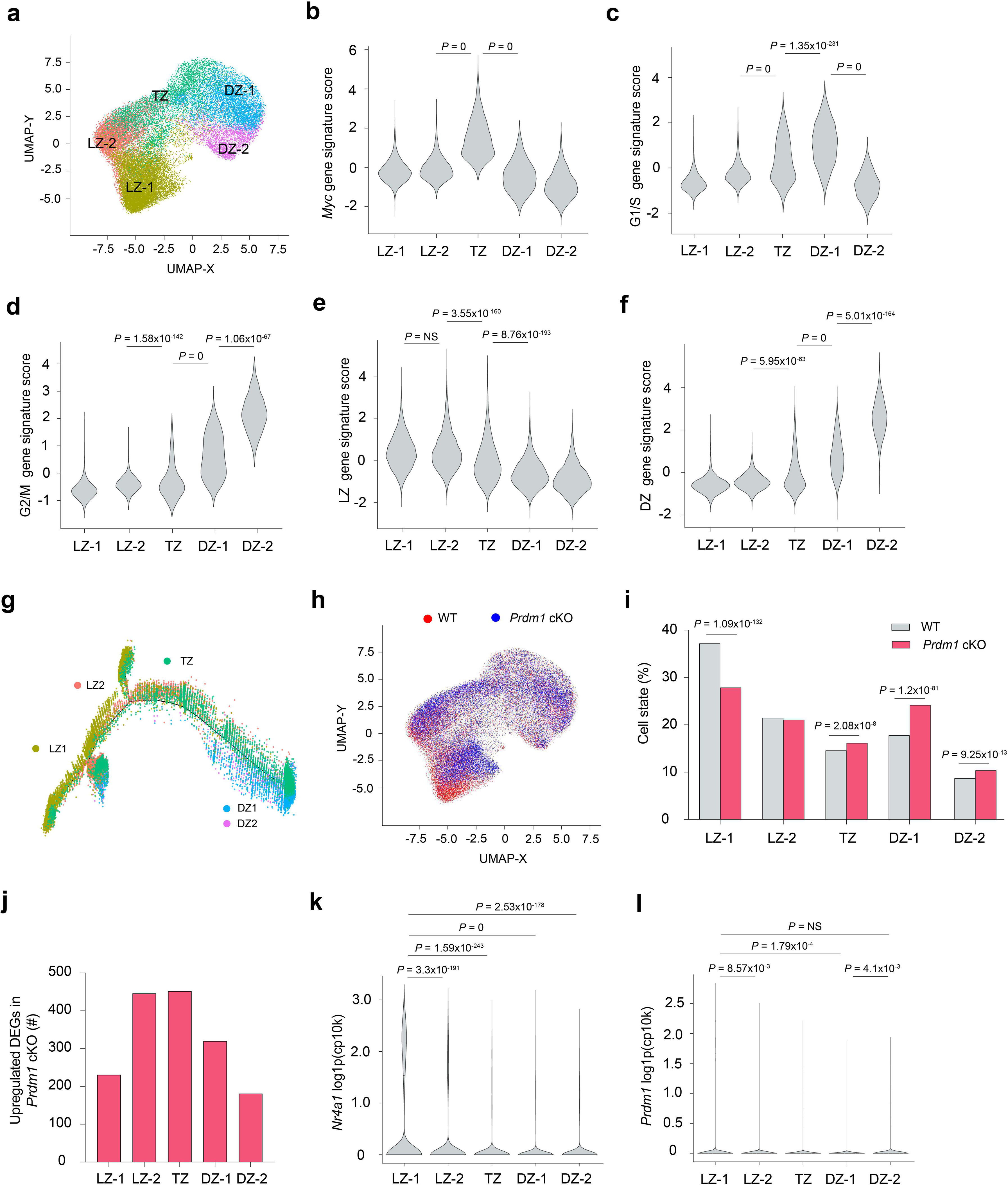

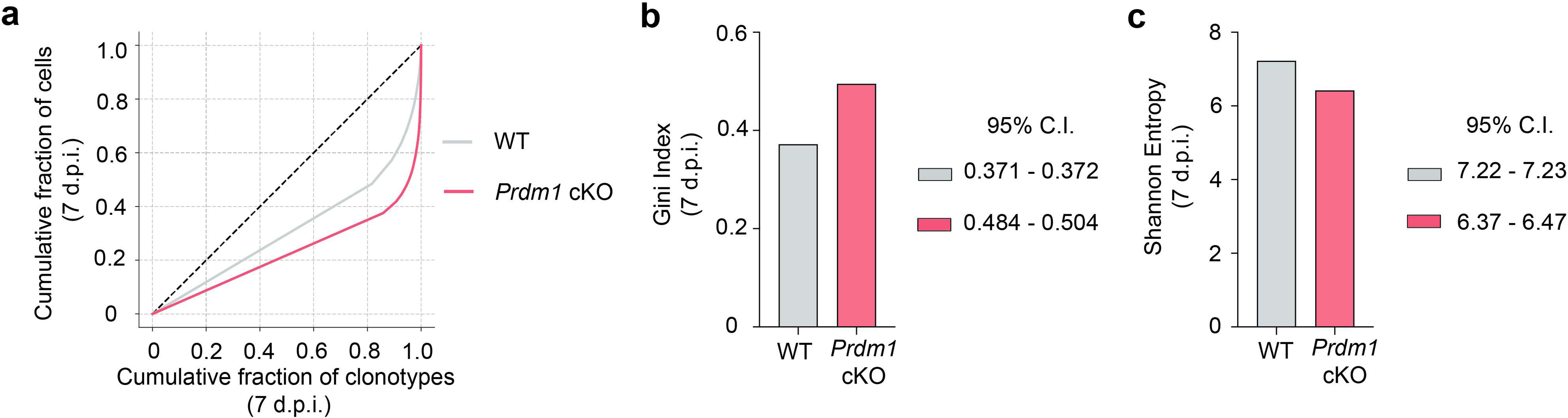

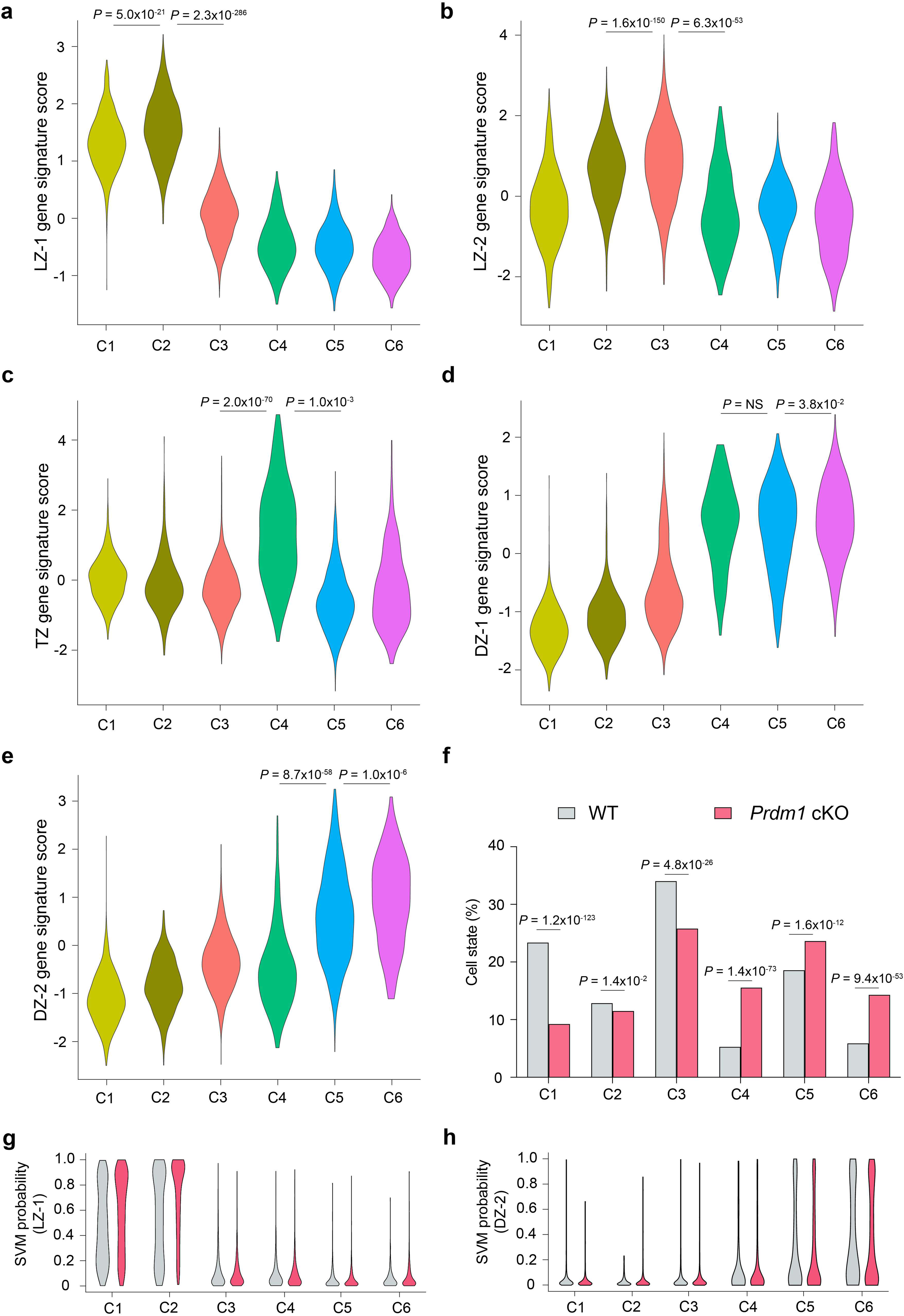

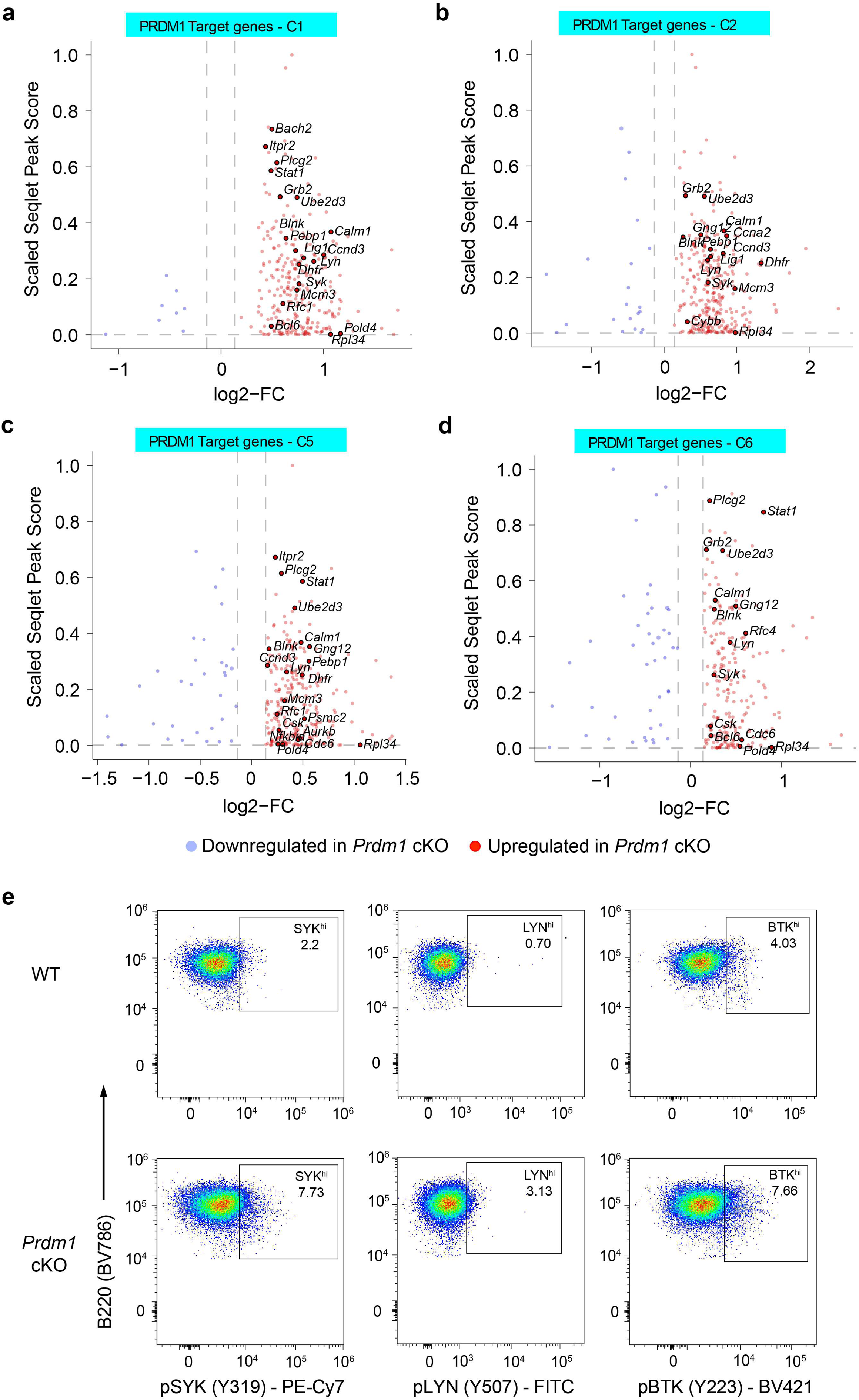

